# Gattaca: Base pair resolution mutation tracking for somatic evolution studies using agent-based models

**DOI:** 10.1101/2021.11.08.467784

**Authors:** Ryan O Schenck, Gabriel Brosula, Jeffrey West, Simon Leedham, Darryl Shibata, Alexander RA Anderson

## Abstract

Gattaca provides the first base-pair resolution artificial genomes for tracking somatic mutations within agent based modeling. Through the incorporation of human reference genomes, mutational context, sequence coverage/error information Gattaca is able to realistically provide comparable sequence data for *in-silico* comparative evolution studies with human somatic evolution studies. This user-friendly method, incorporated into each *in-silico* cell, allows us to fully capture somatic mutation spectra and evolution.

## Introduction

Recent studies examining histologically normal human and murine tissue has shown that a surprising admixture of somatic mutations can exist and even expand to a significant clonal area^1–5^. These studies have largely focused on mutation characteri-zation and have had limited tools to offer explanations for the dynamics driving observed evolutionary trajectories, with only few notable exceptions. Fewer still have begun incorporating agent based models as a tool to explore somatic evolution in spatially constrained tissue^5–8^. Historically, genomes within agent based models have been represented as simple counters or as binary arrays. These studies have lacked the ability to compare, at base pair resolution the mutation spectra or utilize common tools designed for genotypical data (such as dN/dS). Gattaca is the first to provide a means of tracking base pair resolution SNVs within an agent based modeling framework. This cruically provides an ability to accurately capture mutation data on a level comparable to sequencing experiments from the clinic or research settings.

## Implementation

Gattaca is a provided as an easily executed python script consisting of three parts: initialization, execution, and analysis (Figure 1). After the initial setup Gattaca produces a java file that allows for integration into an agent based model (ABM). The only pre-requisite for Gattaca is an installation of snpEff^9^, which provides the necessary base pair resolution reference genome and the tools to access it before Gattaca digests this information for downstream uses.

**Figure 1.**
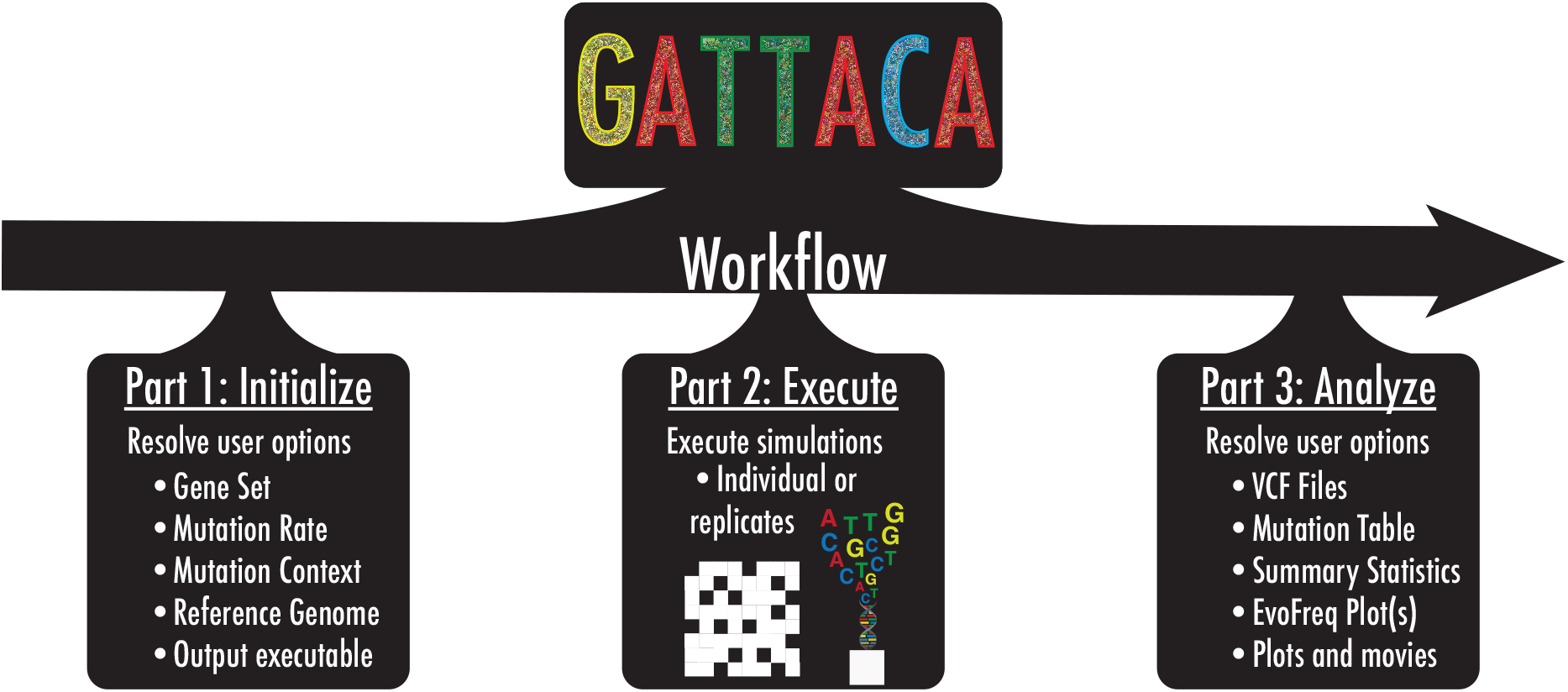
Gattaca is a three part workflow for simulating base pair resolution mutations within the human genome for somatic evolution *in silico* studies. Gattaca consists of three parts, (i) user defines options (initialize), (ii) generate a java executable class for *in silico* simulations with base pair resolution mutation tracking (execution), (iii) analyze the output of these simulations for downstream analysis (analyze).

### Part 1: Initialization

The setup resolves user inputs that includes mutation rates, mutation context probabilities, a gene set, and reference genome choice. Once resolved Gattaca extracts the gene locations from within the users reference genome, a Browser Extensible Data (BED) file is created that snpEff uses to extract bases for each gene. Gattaca then reads a provided mutational context file, in the event of none being provided a uniform probability is used. This file represents the probability of observing a mutation given from the 96 possible mutations within their trinucleotide contexts. Lastly, the mutation rates are scaled to the desired mean mutation rate. The mutation rates are adjusted from the gene specific mutation rates derived from a pan-cancer study^10^. This information is then prepared to generate a Gattaca java class tailored for execution within a HAL ABM^11^, although any ABM framework could be used.

The heart of Gattaca is its ability to track mutations within simulations at a base pair resolution. This requires a series of steps during each cell division where a user checks for mutation. The expected number of mutations per division is given for each gene (*g*_*i*_) by the product of its individual mutation rate *μ*_*gi*_ and its length *L*_*gi*_. Within each mutation check during division a Poisson distribution is used to determine the number of mutations accrued for each gene (*X*_*g*_), so that *X*_*g*_ ∼ *Poisson*(*μ*_*g*_ *\*L*_*g*_).

Determining the specific base that acquires a mutation is based on a multinomial of the 32 possible mutation positions based on trinucleotide contexts. This is drawn from a multinomial distribution based on the 32 possible positions. Once the trinucleotide is determined the base mutation is determined using the mutation context probabilities to determine the mutation type.

### Part 2: Execution

Simulations utilizing Gattaca require the two files that are output by the Gattaca initialization step. These files, a java Gattaca class and a csv file with loci information, will be placed within the scope of your executable HAL model^11^. Details on using HAL can be found at (http://halloworld.org). Once these are added to HAL, the Gattaca class will require initialization for a founding clone/population. Gattaca ties conveniently into the HAL phylogeny tracker requiring minimal additional computational overhead. Once Gattaca is initialized a function call to *_RunPossibleMutation* will be required during each division that will trigger the possibility of mutation upon division as outlined above. A detailed tutorial on integrating Gattaca and HAL can be found at https://github.com/MathOnco/Gattaca.

### Part 3: Analysis

Once simulations are complete Gattaca introduces the appropriate noise for each mutation type, one of two ways (adapted from^12^). The true variant allele frequency (assuming heterozygosity), *VAF*_*t*_, is given from 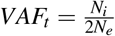, where *N*_*i*_ is the number of cells with a given muation and *N*_*e*_ is the population size. The user can provide a list of depths for mutations within an experimental cohort or define a single value *sequencing* depth. If the user sets a single value for depth (*d*) the number of reads calculated for the depth of a variant, *D*_*i*_, is drawn from a Poisson distribution, which yields *D*_*i*_ ∼ *Poisson*(*d*). If a user provides a distribution of depths from an experimental cohort Gattaca determines the shape parameters (*k*_*c*_ and *p*_*c*_) defining a gamma distribution to obtain *D*_*i*_ so that *D*_*i*_ = *Gamma*(*k* = *k*_*c*_, *p* = *p*_*c*_). The number of reads for a given variant (*f*_*i*_) is finally determined by *f*_*i*_ = *B*_0_(*n* = *D*_*i*_, *p* = *VAF*_*t*_). By taking the sequenced VAF 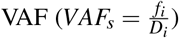 and applying a threshold (typically 0.005 to 0.1 depending on sequencing depth) Gattaca yields mutations that are comparable to what may be observed from sequencing of tissue.

Once the variants are called based on the corrected VAF, variants are annotated with snpEFF and mutational position information is obtained. The user can output this information as a mutational table for every desired timepoint and every replicate simulation. As an additional output option users can also export variants from their simulations as a variant call format (VCF) file. This option allows for easy use in several bioinformatics downstream tools. Lastly, the execution of the analysis component of Gattaca provides several summary statistics for evolutionary dynamics, such as 1*/ f* ^12^, first incomplete moment^3,13^, an EvoFreq plot^14^, and a crude dN/dS measurement. We note that a true dN/dS would be expected to be the same across all simulations unless the user implements functional heterogeneity within their simulations based on a single, or collection of, point mutations.

### Case Study 1: Dimensionality

Gattaca allows us to track base pair resolution genomes across any agent based modeling dimension. Recent interest by ourselves and others in understanding how spatial architecture may affect clonal dynamics and measurements of neutrality motivates our case study^1,3,6,15,16^.

Here we have constructed two simple agent based models (ABM) of cell turnover in three different dimensions, zero-(0*D*), two-(2*D*), and three-dimension (3*D*), to showcase and compare the mutational profiles and clonal dynamics that Gattaca allows its users to evaluate. In addition, we perform the simulations for these three dimensions and two model types for three different total final population sizes to demonstrate the functionalities and outputs of Gattaca (Figure 2). The two model types differ only in the number of cells that are present at initialization. The fully seeded model initializes by placing an agent with its unique genome at every lattice point or until the carrying capacity is reached in the 0*D* case. The second simple ABM is initialized only with a single cell at a random position within the simulated domain, or simply a population size of one for the 0*D* case (Figure 2). These two simple model types can be conceptualized as a naive tissue type of model to compare with a stem cell growth model similar to the idea that cancer originates from a single transformed clone. Here we introduce no functional heterogeneity across the different genomes that emerge through mutation at each timepoint governed by the conditions set in Gattaca.

**Figure 2.**
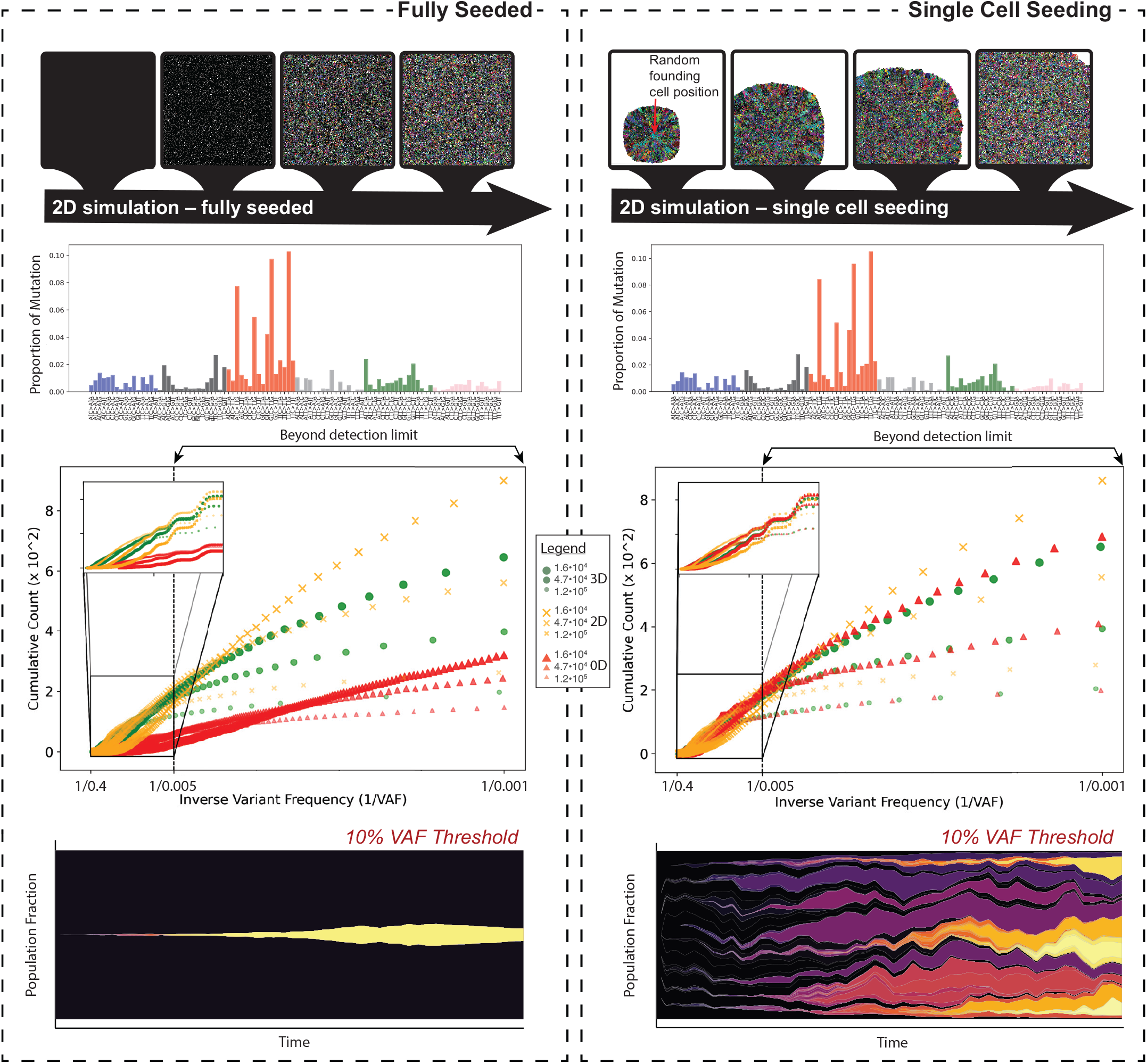
Results comparing the fully seeded (*left*) and single cell seeding (*right*) model types and their corresponding mutation profiles and clonal dynamics. For each case, the mutation proportion across the 96 mutation trinucleotides is shown for one of the 2*D* simulation replicates. The 1*/ f* values for each of the modeled dimensions is shown for three different population sizes, the inset shows the 1*/ f* distribution for that which would be within the limits of detection (a generous 0.005 VAF at high depths). Beneath this, the same 2*D* replicate that is shown in the mutation spectrum plot is used to highlight the differences in clonal dynamics using an EvoFreq plot with a 10% VAF cutoff between fully seeded and single cell seeding.

Within the two models we use the same parameters so as to be able to more accurately compare across the different dimensions. Each model across all dimensions uses the same birth/death function. The birth rate (*λ, λ* = 0.4) is scaled by the carrying capacity (*k*) and population size (*N*_*T*_) at every time point of either the domain (e.g. number of lattice points) or as a set parameter in the 0*D* case. The equation governing this scaled birth rate (*λ*_*T*_) is given by 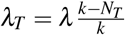. If a random number ([0, 1]) is less than the death parameter (*ρ*) plus *λ*_*T*_ a death or birth may happen for a given cell. The probability of a birth event given an empty lattice position (2*D* and 3*D* only) is given by *P*(*Birth*) = *ρ* + *λ*_*T*_. If a random number ([0, 1]) is less than this birth event value a cell will die, if not the cell is able to divide.

When initializing Gattaca for these simulations an overall mutation rate of 3.2 *\** 10^*−*^9 was used and the mutation spectrum defined was given from a sampled cohort of Large B-Cell Lymphoma whole exome sequencing (this is available in the gattaca example code). When we analyze these mutation spectrums, post simulation we observe similar distributions of mutation types across all dimensions and model types consistent with mutation processes expected, based on the Gattaca initialization (Figure 2 mutation spectrums). The differences that are observed largely depend upon the dimensionality of the model chosen and the tissue type modeled. In the cases where the domain (or carrying capacity for 0*D*) is fully seeded we see that the 1*/ f* distributions of variant allele frequencies is similar in the 3D and 2D cases (Figure 2). Contrasting this with the single cell seeding case we see that the 0*D* and 3*D* cases are the most similar while 2*D* appears to reveal a different distribution (Figure 2). These results suggest that the modeling dimension is an important consideration for the research question. As expected most of the clones that are obsered are below the detection limits of common methodologies, but can be captured here. The clonal dynamics, as demonstrated by the EvoFreq plots^14^ illustrates that spatially constrained clones competing with one another are rarely able to expand beyond 10% VAF in the fully seeded cases while several clones reach this size during simulations with single cell seeding.

### Case Study 2: Wounding

Within the first case study we utilized Gattaca across two different types of models and three different dimensions. Next we wanted to evaluate if wounding within these models would alter the observed clonal dynamics as the spatial constraints for certain clones is relaxed when cells are removed in a wounding event (Figure 3A). In all simulations, each ABM is seeded by a single cell. Wounding begins once the thousandth timestep is reached (Figure 3B). After this, wounding occurs at time steps where the population is greater or equal to 85% of the total possible population (as dictated by the domain size). For the 2*D* and 3*D* simulations cells are killed by wounding within a circular and spherical manner, respectively. The number of cells killed through each wounding event is kept similar by adjusting the radius between 2*D* and 3*D* simulations, while in the 0*D* case, the number of cells killed is an equivalent number of cells. The same birth/death dynamics and equations used in case study one are used here, because the probability of birth is modulated by the number of cells (*i*.*e*. the probability of birth is reduced as the carrying capacity of the system is reached) a wounding event acts to increase cell divisions where empty sites are present and thus allows clones to expand into the wounded areas.

**Figure 3.**
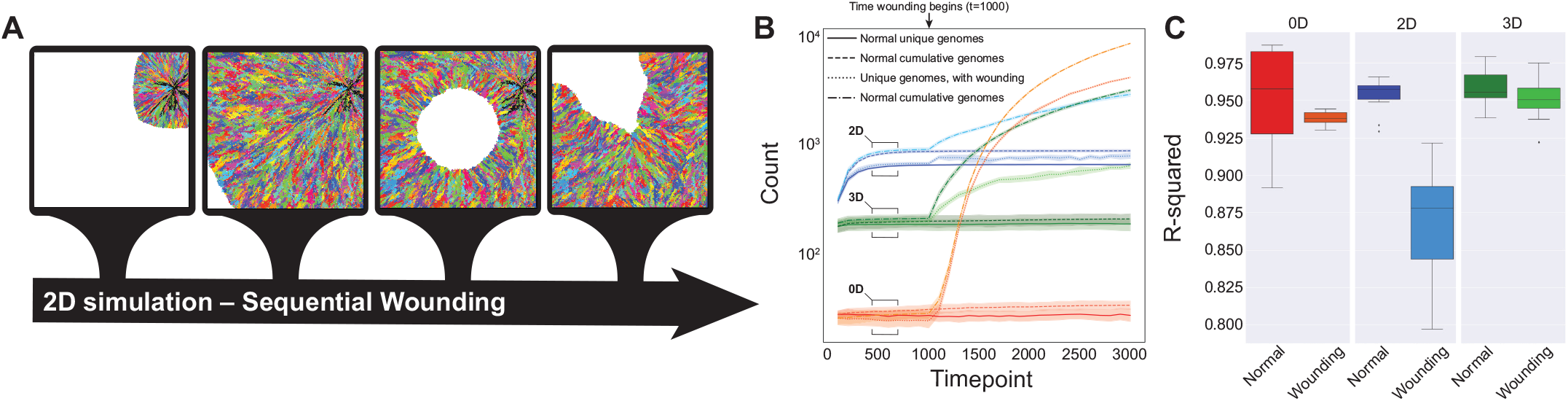
Illustration of the repeated wounding of the single cell model in 2*D* where colors represent clones that differ by at least one mutation (A). For each of the dimensions the cumulative and unique genomes is given over the course of simulations (B). R-squared values for the linear regression on 1*/ f* distributions for mutations is plotted for all dimensions with and without wounding (C) for all replicate simulations.

When we examine the differences between the wounding and non-wounding simulation’s cumulative and unique genomes over time we see a clear signal at the time wounding occurs. At this point, space is open and rapid cell proliferation refills the areas where the wound occurred (in 0*D* this results in rapid proliferation back to carrying capacity). As cells divide and mutate a large number of unique genomes appear over time (Figure 3B). We see that the number of unique genomes in the 0*D* case increases drastically faster than those in the 2*D* and 3*D* cases, this is due to the mechanism where clones in the 0*D* case are chosen at random to be killed while whole or near whole subclonal populations are removed in the 2*D* and 3*D* simulations. Interestingly, when we compare the 1*/ f* distributions through their R-squared values, from linear regression analysis, we see that in the 2*D* wounding case the relaxation of spatial constraints appears to drive a signal of non-neutral dynamics in a system that is functionally homogeneous where slight fitness advantages are conferred through room to expand (Figure 3C).

## Conclusions

Here we have presented Gattaca, the first base pair resolution mutation tracking *in silico* genome for agent based modeling. Gattaca provides a powerful tool to track mutations through time and space to compare with patient and murine samples. We have demonstrated this by comparing the genomes and clonal dynamics that Gattaca provides across different modeling dimensions and model choices. We then show through a second use case that wounding can show evidence of selection, but only in the 2*D* wounding case. This sets an important precedent that modeling choices around dimensionality can significantly impact the measures of neutrality.

Gattaca provides a highly customizable framework that is easily implemented into users agent based simulations for evaluating somatic evolution in normal or disease tissue. Through the incorporation of common bioinformatics and genotypic outputs (variant call format files) used frequently in clinical and experimental approaches users can quickly analyze and compare mutation spectra, burden, heterogeneity, and selection between their samples and *in silico* models.

## Code availability

Gattaca is available through GitHub (https://github.com/MathOnco/Gattaca). There is a read me available within the GitHub repository with further instructions on how to utilize Gattaca.

## Acknowledgements

ROS is supported by the Wellcome Trust (grant no. 108861/7/15/7) and the Wellcome Centre for Human Genetics (grant no. 203141/7/16/7). ROS and ARAA are supported by the Cancer Systems Biology Consortium grant from the National Cancer Institute (grant no. U01CA23238) and the Moffitt Cancer Center of Excellence for Evolutionary Therapy. SL is supported by the Wellcome Trust (grant no. 206314/Z/17/Z). DS is supported by the Cancer Systems Biology Consortium grant from the National Cancer Institute (grant no. U54CA217376 and grant no. P01 CA196569).

## Author contributions statement

RS, DS, and ARAA conceived the idea Gattaca for ABM. Case studies one and two were done by RS and GB with assistance from JW. The manuscript was written by RS, GB, JW, and ARAA. All authors reviewed the manuscript.

